# HNF4α mediated QPRT expression in the kidney facilitates resilience against acute kidney injury

**DOI:** 10.1101/2023.05.23.541987

**Authors:** Amanda J. Clark, Marie Christelle Saade, Vamsidhara Vemireddy, Kyle Q. Vu, Brenda Mendoza Flores, Valerie Etzrodt, Erin J. Ciampa, Huihui Huang, Ayumi Takakura, Kambiz Zandi-Nejad, Zsuzsanna K. Zsengellér, Samir M. Parikh

## Abstract

Nicotinamide adenine dinucleotide (NAD+) levels decline in experimental models of acute kidney injury (AKI). Attenuated enzymatic conversion of tryptophan to NAD+ in tubular epithelium may contribute to adverse cellular and physiological outcomes. Mechanisms underlying defense of tryptophan-dependent NAD+ production are incompletely understood. Here we show that regulation of a bottleneck enzyme in this pathway, quinolinate phosphoribosyltransferase (QPRT) may contribute to kidney resilience. Expression of QPRT declined in two unrelated models of AKI. Haploinsufficient mice developed worse outcomes compared to littermate controls whereas novel, conditional gain-of-function mice were protected from injury. Applying these findings, we then identified hepatocyte nuclear factor 4 alpha (HNF4α) as a candidate transcription factor regulating QPRT expression downstream of the mitochondrial biogenesis regulator and NAD+ biosynthesis inducer PPARgamma coactivator-1-alpha (PGC1α). This was verified by chromatin immunoprecipitation. A PGC1a-HNF4α -QPRT axis controlled NAD+ levels across cellular compartments and modulated cellular ATP. These results propose that tryptophan-dependent NAD+ biosynthesis via QPRT and induced by HNF4α may be a critical determinant of kidney resilience to noxious stressors.

## INTRODUCTION

Acute kidney injury (AKI) affects roughly 20-25% of hospitalized children and adults and is associated with substantial morbidity and mortality.^1, 2^ Normal renal tubule function requires efficient adenosine triphosphate (ATP) generation, and impairment in renal energy metabolism is a key feature of AKI.^3^ Nicotinamide adenine dinucleotide (NAD+) is a critical electron receptor required for ATP production. Reduction of NAD+ becomes rate-limiting for ATP production.^4^ During AKI, degradation of cellular NAD+ is upregulated and, maladaptively, NAD+ biosynthesis is attenuated.^5–9^ Suppression of NAD+ biosynthesis, particularly the de novo pathway of NAD+ biosynthesis from tryptophan, has been described as an element of AKI in experimental models and humans.^7,8,10–12^ Specifically, the bottleneck enzyme of de novo NAD+ biosynthesis, quinolinate phosphoribosyltransferase (QPRT), is suppressed in post-ischemic kidney injury.^7^ Despite existence of alternate NAD+ biosynthetic pathways, QPRT+/- mice exhibit reduced basal kidney NAD+, reduced basal kidney ATP, and increased susceptibility to post-ischemic AKI.^7^

We have previously shown that QPRT is regulated by PPARgamma coactivator 1 alpha (PGC1α), a transcriptional co-activator known to regulate diverse genes involved in energy metabolism, mitochondrial biogenesis, and mitochondrial quality control.^13–18^ PGC1α is suppressed in renal injury with resultant decreases in NAD+ and ATP; furthermore, reduced PGC1α exacerbates AKI.^13–17^ Conversely, induction of PGC1α protects mice from post-ischemia injury, cisplatin injury, and experimental renal fibrosis.^13, 14, 18, 19^ A transcription factor linking PGC1α to NAD+ biosynthesis has yet to be identified.

Hepatocyte nuclear factor 4 alpha (HNF4α) is a nuclear receptor and transcription factor that regulates genes associated with lipid metabolism, glucose metabolism, and cellular differentiation. In the liver, HNF4α modulates insulin responsiveness, with HNF4α loss of function mutations leading to mature onset diabetes of the young (MODY) Impaired HNF4α activity has also been linked to non-alcoholic fatty liver disease,^20, 21^ inflammation,^22^ fibrosis, and cirrhosis.^23^ HNF4α is also critical for hepatocyte differentiation from stem lineages.^24^ In the developing kidney, HNF4α is necessary for proximal tubular cell differentiation from progenitor cells.^25^ In the adult kidney, HNF4α expression and enhancer binding is decreased in acute injury.^26, 27^ Re-expression of HNF4α appears to be a marker and requirement of cellular repair.^27, 28^ However, little is known about HNF4α-mediated metabolism in the healthy kidney or its role in kidney cellular injury.

We hypothesized that de novo NAD+ biosynthesis -- the tryptophan pathway of NAD+ biosynthesis -- suppression would be a consistent feature of toxic AKI. We further hypothesized that augmented kidney QPRT expression would be sufficient to protect against toxic insults. Finally, we tested whether HNF4α is an essential transcription factor that regulates renal NAD+ biosynthesis via QPRT modulation, thus implicating HNF4α in metabolic resilience against AKI.

## METHODS

### Mouse Models

All experiments were conducted in compliance with the NIH Guide for the Care and Use of Laboratory Animals and approved by an institutional animal care and use committee (IACUC). QPRT +/- mice were previously described.^7^ iNephQPRT mice were generated by crossing a Pax8-rtTA mice (Jax #007176)^29^ to a custom tetO-QPRT mouse created via pronuclear microinjection of a TetO-QPRT plasmid on a pBT264 backbone into C57bl/6J zygotes. Genotyping was performed using Transnetyx. Experimental animals were bred to be hemizygous for both the Pax8-rtTA and the tetO-QPRT transgenes. Siblings carrying only the tetO-QPRT transgene were used as controls. Overexpression was induced via delivery of doxycycline (Fisher Scientific) in the water (0.2mg/mL+ 5% dextrose)^30^ or chow (Teklad 625mg/kg) offered ad lib. Doxycycline was started at the time of injury and continued until sacrifice for both iNephQPRT mice and controls. iNephPGC1α (Pax8-rtTA, tetO-PGC1α) mice were previously described.^14^ Cisplatin nephrotoxicity was induced via a single intraperitoneal (IP) injection of cisplatin (Aldrich) 25mg/kg dissolved in normal saline, and animals were harvested at 72 hours. Folic acid AKI was generated by single IP injection of folic acid (Fisher) 250mg/kg dissolved in 0.3M sodium bicarbonate, and animals were harvested at 24 hours. Only 8--12-week-old male mice were used as trials in females resulted in excessive mortality.

### Metabolomics

Metabolomics were performed on whole kidney lysate performed by Metabolon. In brief, proteins were precipitated with methanol followed by centrifugation. The resulting extract was divided into five fractions: two for analysis by two separate reverse phase (RP)/UPLC-MS/MS methods with positive ion mode electrospray ionization (ESI), one for analysis by RP/UPLC-MS/MS with negative ion mode ESI, one for analysis by HILIC/UPLC-MS/MS with negative ion mode ESI, and one sample was reserved for backup. A pooled matrix sample served as a technical replication, extracted water samples served as blanks, and QC standards were spiked into every sample. Raw data was extracted, peak-identified, and QC processed using Metabolon’s hardware and software. Compounds were identified by comparison to a maintained library of known compounds based on retention time/ index, mass to charge ratio, and chromatographic data. Peaks were quantified using area under the cure.

### RT-qPCR and biochemical assays

For cell and animal experiments, RNA was isolated from cell of tissue lysate using Bio-Rad Aurum Total RNA Mini Kit. cDNA was made using Bio-rad iScript Advanced cDNA Synthesis Kit. qPCR was performed using SYBR green, and the listed primers (**Supplementary Table S1**). BUN was measured using a colorimetric assay (Invitrogen). Serum creatinine was measured using capillary electrophoresis at the University of Texas Southwestern renal physiology core. ATP was measured using a luminescent assay (Abcam).

### Histopathology

Kidneys were placed in formalin at the time of harvest. Before staining, kidneys were rinsed, dehydrated, cleared, and paraffin embedded. Serial sections were obtained by rotary microtome at 5um thickness. PAS staining was performed manually by the Research Histo Pathology Core at the University of Texas Southwestern. In detail, slides were deparaffinized, run to water, and bound carbohydrates were impregnated with 0.5% periodic acid. Subsequently, slides were rinsed in water and developed in Schiff’s reagent (Sigma). Excess Schiff’s was removed with water and hematoxylin nuclear counterstain was applied followed by water rinse and bluing. Slides were then dehydrated, cleared, and cover-slipped with synthetic resin. Injury scoring was performed by a blinded pathologist or by two blinded physicians and averaged. Tubular injury was defined as denudation of the renal tubular cells, loss of brush border, tubular dilation, intratubular casts, and cell sloughing with the following scoring system, 0 = no evidence, 1 = <10% involved, 2 = 10-25% involved, 3 = 25-50% involved, 4 = 50-75% involved, and 5 = >75%involved.

### Cell models

Unless otherwise specified cell experiments were conducted in HK2 (ATCC), an immortalized human proximal tubule cell line at an early passage grown in DMEM with 10% fetal bovine serum (FBS). PGC1α or HNF4α overexpression was induced using plasmids from Addgene as gifts from Toren Finkel (#10974)^31^ and Gerhart Ryffel (#31100).^32^ QPRT plasmid was purchased from Origene (RC202960). Plasmids were transfected using Lipofectamine 3000 (Invitrogen). siRNA knockdowns were performed using pre-made siRNA smartpools (Horizon) and transfected with siRNAiMax (Invitrogen). Supplemental studies were conducted in immortalized pig proximal tubule cells (LLC-PK1) (ATCC) grown in Medium 199 with 3% FBS. A stable PGC1α cell line was created by infecting LLC-PK1 cells with a previously published lentivirus^18^ followed by selection with puromycin.

### NAD+ Biosensors

Biosensor plasmids were a gift from Dr. Richard Goodman.^33^ Sensors were transfected into cells using Lipofectamine 3000. Cells with sensors were imaged using confocal microscopy at both an NAD+ responsive 488nm excitation and a non-NAD+ responsive internal control that excited at 405nm. Relative NAD+ was quantified using Image J software to calculate the ratio of 488 fluorescence to 405 fluorescence per cell. Values were normalized to empty vector control cells. Results are displayed as the inverse of the fluorescence calculation so that higher values correlate with increased NAD+ abundance.

### ChIP qPCR

Chromatin immunoprecipitation (ChIP) was performed on kidney lysate from iNephPGC1α mice using the EpiQuik Tissue Chromatin Immunoprecipitation Kit. Tissue was crosslinked in 1% formaldehyde and mechanically disrupted using a cell strainer. Cells were lysed and chromatin was sheared using sonication. Immunoprecipitation was accomplished using plate based-beads in the EpiQuik Kit along with HNF4α antibodies (Novus NB100-1783 and Abcam ab181604) to validate enrichment at *QPRT*. Comparison between iNephPGC1α and control kidneys used Abcam antibody alone. Anti-goat non-immune IgG was used as a non-targeting control. QPRT qPCR was completed using SYBR green mouse *QPRT* gDNA primers (**Supplemental TableS1**).

## RESULTS

In a cisplatin model of AKI, unbiased metabolic profiling of injured and control kidney tissue identified accumulation of de novo NAD+ biosynthetic intermediates, reduction of NAD+ itself, and reduction of metabolites arising from NAD+ degradation (**Fig1 A-C**). Semiquantitative qPCR confirmed suppression of NAD+ biosynthesis enzymes (**Fig 1D**) in cisplatin-treated kidneys. Among these, the suppression of QPRT was the most marked. Given the heterogeneity of functional impairment in the cisplatin model, we evaluated the proportionality of QPRT expression with injury severity. Indeed, QPRT was proportionally suppressed to injury severity as quantified by several readouts: LCN2 kidney mRNA, BUN, and serum creatinine as injury markers (**Fig 1E-F**).

**Figure 1:**
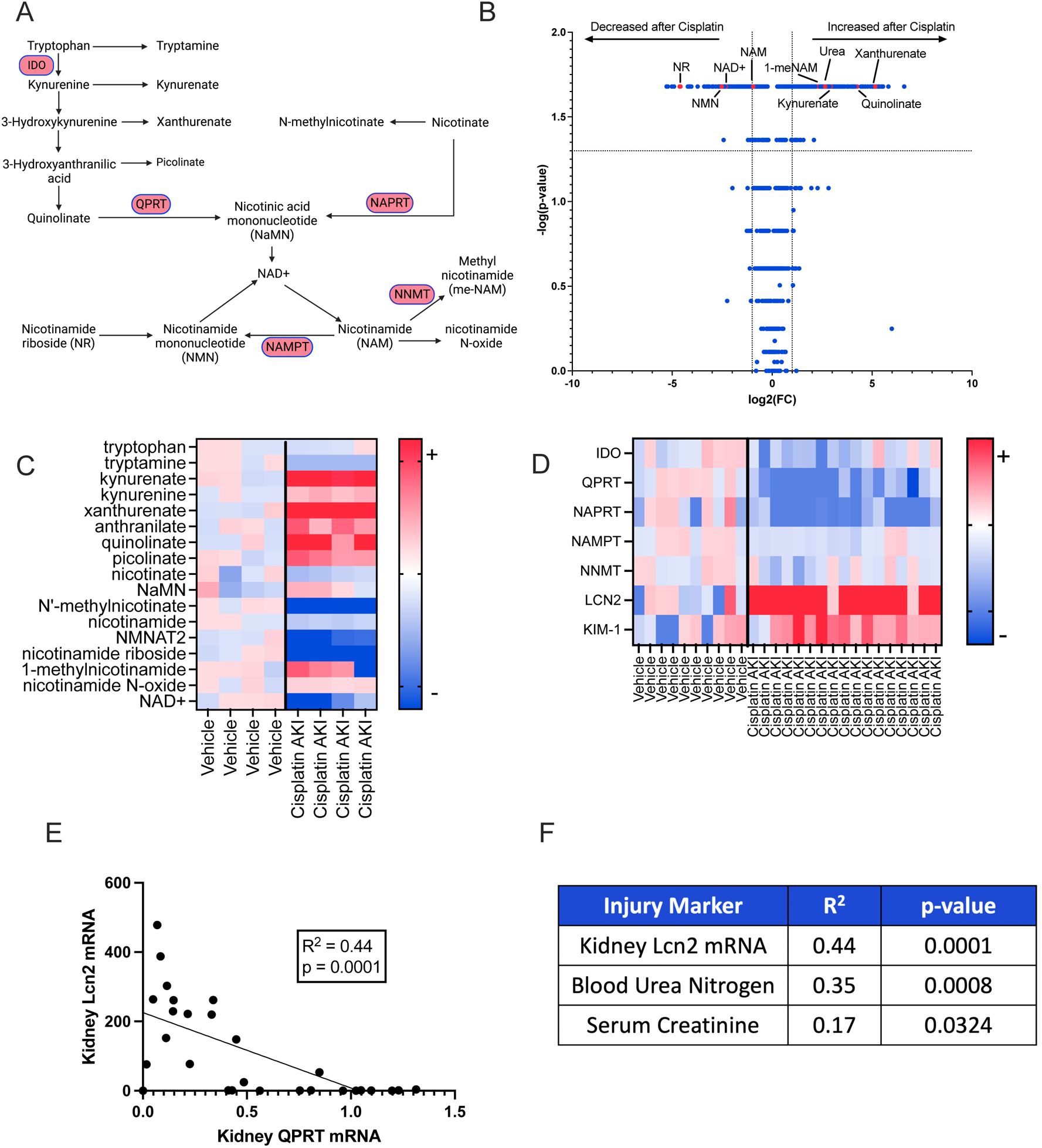
De novo NAD+ Biosynthesis suppression is a component of nephrotoxic AKI. **A.** Schema of NAD+ Biosynthesis. **B.** Volcano plot of mouse kidney metabolites from mice that received cisplatin or vehicle. P-values calculated using Mann-Whitney. Dotted lines indicate p value greater of 0.05 (horizontal) and fold change < or > 2 (vertical). **C.** Heat map showing individual values of selected metabolites. Values are normalized to average of vehicle animals and presented as fold change. Darkest colors indicate fold change ≥6 (red) or ≤-6 (blue). **D.** Heatmap representing gene expression of selected genes in WT male mice who received 25mg/kg IP cisplatin (n=17) or Vehicle (n=10). Data are normalized to the vehicle average and expressed as fold change. Darkest colors indicate fold change ≥5 (red) or ≤-5 (blue). **E.** Correlation of kidney QPRT expression to kidney Lcn2 expression in WT mice that received cisplatin or vehicle. Correlation and p-value calculation using a simple linear regression, **F.** Correlation of 3 kidney injury markers to kidney QPRT mRNA expression. Correlation and p-values calculated with simple linear regression.

Because intracellular levels of NAD+ are distinct across cellular compartments with compartment-specific actions,^33^ we sought to further understand the metabolic implications of QPRT suppression by assessing which cellular compartments demonstrated NAD+ modulation following manipulation of QPRT expression. We first established that QPRT plasmid or siQPRT could be transfected into cells to overexpress or suppress QPRT expression, respectively (**Supplemental FigS1A-B**). We then employed compartmentalized NAD+ biosensors^33^ to elucidate the effect of QPRT modulation on compartment-specific NAD+ levels (**Fig 2A**). QPRT siRNA decreased NAD+ in all cellular compartments while QPRT overexpression increased NAD+ in all cellular compartments. (**Fig 2B-G**). QPRT siRNA reduced cellular ATP (**Fig 2H**) whereas QPRT overexpression increased ATP (**Fig 2I**) relative to control cells. This pattern mirrored PGC1α overexpression, which also increased NAD+ in all cellular compartments and increased cellular ATP (**Supplemental FigS2A-E**)

**Figure 2:**
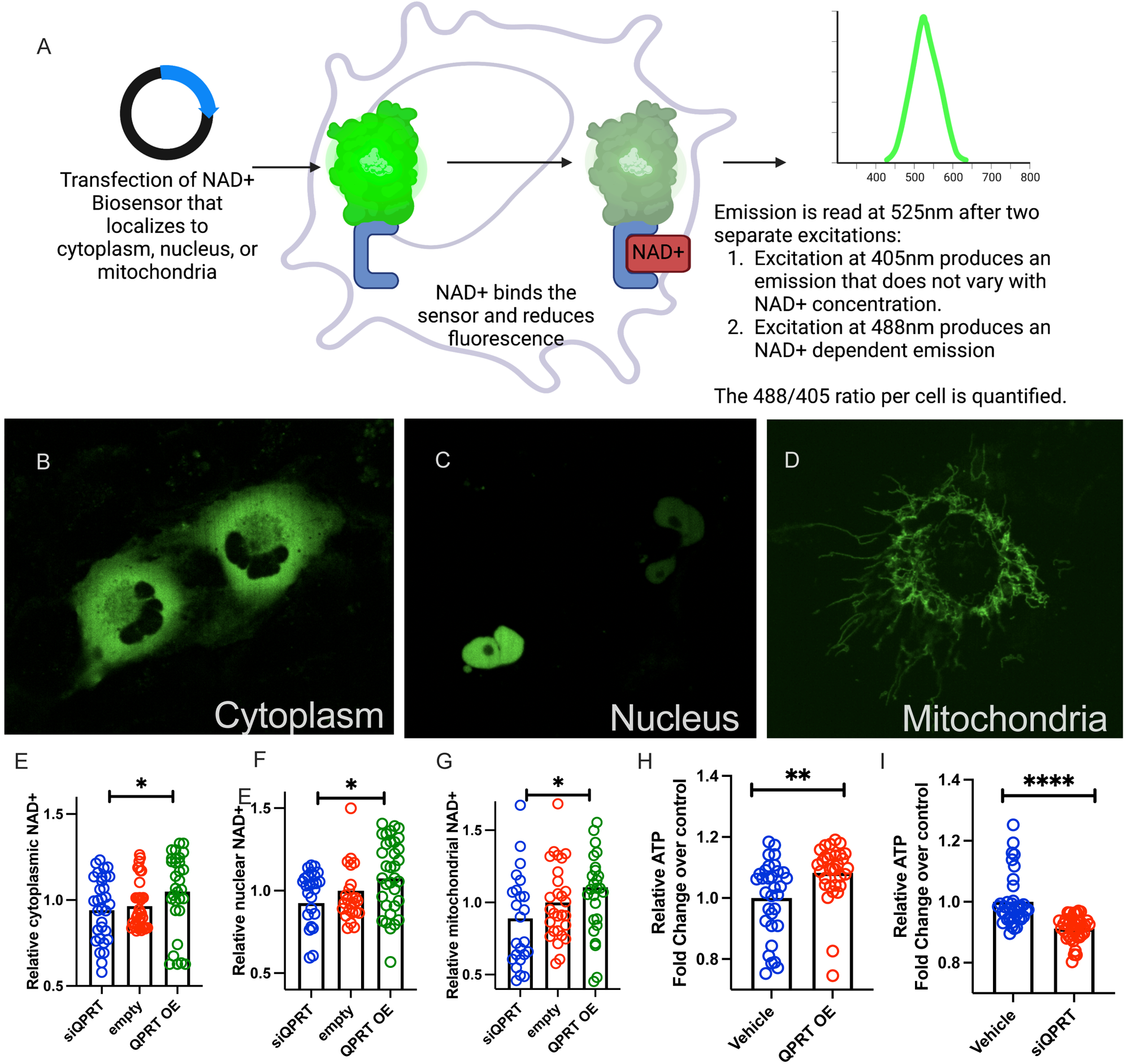
QPRT expression mediates NAD+ and ATP. **A.** Schematic describing a compartment-specific NAD+ biosensor. **B-D** Representative images of compartment-specific NAD+ biosensor in use. Relative cytoplasmic (**E**), nuclear (**F**), and mitochondrial (**G**) NAD+ in QPRT overexpression (OE), control, and siQPRT knock down in HK-2 cells. **H.** Relative ATP in QPRT OE and Control **I.** Relative ATP in siQPRT and Control. P-values were calculated using Mann-Whitney. * = p<0.05, ** = p<0.01, *** = p<0.001, **** = p<0.0001.

We next evaluated the effect of distinct nephrotoxins in vivo. After cisplatin (**Fig 3A**), QPRT +/- mice developed higher serum creatinine (**Fig 3B**), higher BUN (**Fig 3C**), increased expression of renal LCN2 (**Fig 3D**), and worse histological injury (**Fig 3E-F**) compared to wild type littermates. In the unrelated nephrotoxicity that arises 24 hours after systemic folic acid administration (**Fig 3G**), QPRT is also suppressed (**Supplemental FigS3A**), QPRT +/- mice demonstrated a trend toward more elevated serum creatinine (**Fig 3H**), significantly increased BUN (**Fig 3I**), and more severe histological injury (**Fig 3J-K**) compared to littermate controls.

**Figure 3:**
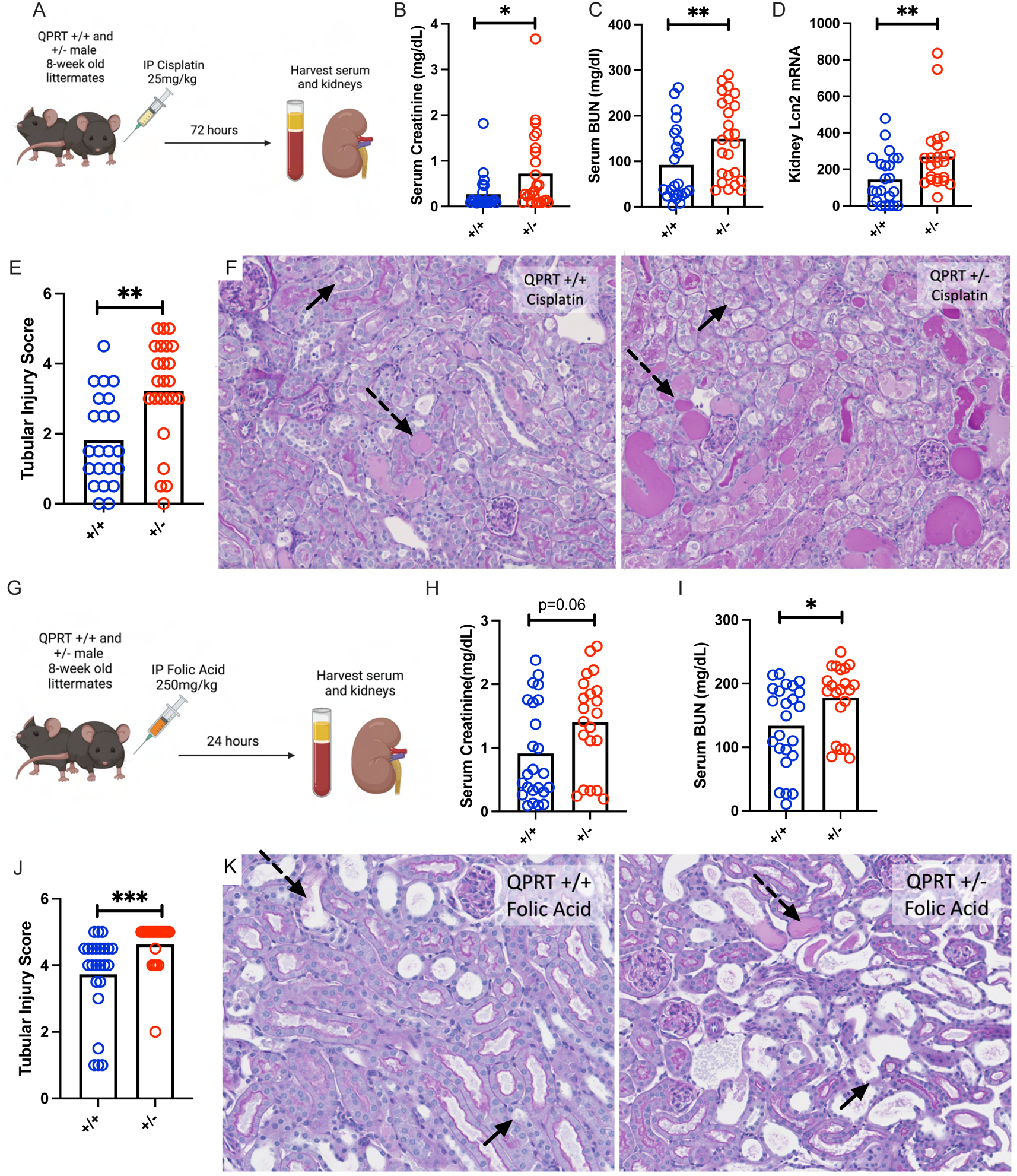
QPRT +/- mice are more susceptible to two distinct models of nephrotoxic AKI. **A.** Schema of Cisplatin AKI experiment. **B.** Serum creatinine, **C.** serum BUN, and **D.** renal mRNA expression of Lcn2 after cisplatin (72 hours after 25mg/kg IP) in QPRT +/+ (n=23) and QPRT +/- (n=25) littermates. All compared with Mann-Whitney. **E.** Tubular injury scoring from histology. F. Representative PAS images after cisplatin. **G.** Schema of folic acid AKI experiment. **H**. Serum creatinine, **I.** serum BUN, and **J.** tubular injury scoring after folic acid (24 hours after 250mg/kg IP) in QPRT +/+ (n=24) and QPRT +/- (n=21) littermates. **K.** Representative PAS images after folic acid. Dotted arrows represent dilated tubules with intratubular casts. Solid arrows represent necrotic cells and denuded tubules. * = p<0.05, ** = p<0.01, *** = p<0.001, **** = p<0.0001.

To investigate whether overcoming QPRT suppression in toxic AKI was sufficient to mitigate AKI, we generated a novel, renal tubule specific, inducible QPRT overexpression mouse (iNephQPRT) by breeding Pax8-rtTA mice to tetO-QPRT knock-in mice (**Fig 4A**). These animals exhibited non-leaky kidney specific overexpression of QPRT after administration of doxycycline (**Supplemental FigS3B**). When given cisplatin (**Fig 4B**), iNephQPRT mice exhibited lower BUN (**Fig 4C**) and less expression of kidney LCN2 mRNA compared to control littermates. While there was no difference in serum creatinine (**Fig 4E**) and only a trend toward worsened histological injury (**Supplemental FigS3C**), we found that the degree of renal injury inversely correlated with QPRT expression (**Fig 4F**). We next evaluated acute folic acid nephropathy in this gain-of-function model (**Fig 4G**). In this model, we used dox chow to induce overexpression to overcome any confounding introduced from the dextrose administered with dox water. Compared to controls, iNephQPRT mice developed less severe injury as measured BUN (**Fig 4H**), serum creatinine (**Fig 4I**), and quantification of histological injury (**Fig 4J-K**).

**Figure 4:**
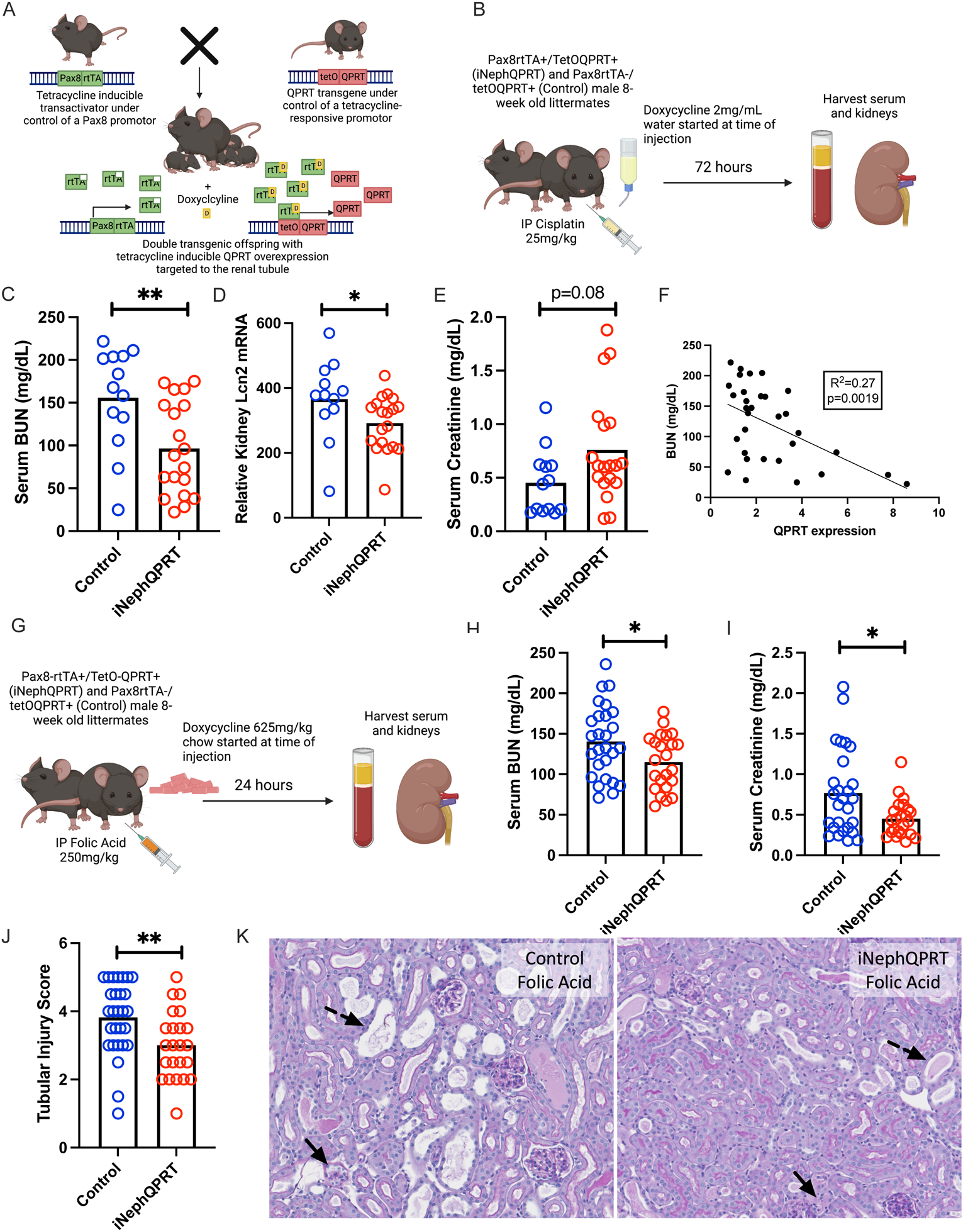
iNephQPRT mice are protected against two distinct models of nephrotoxic AKI. **A.** Schema of iNephQPRT mouse generation. **B.** Schema of iNephQPRT cisplatin experiment. **C.** Serum BUN, renal lcn2 mRNA (**D**), and serum creatinine (**E**) in iNephQPRT (n=19) and control (n=13) littermates after cisplatin (72 hours after cisplatin 25mg/kg IP). **F.** Correlation of serum BUN with QPRT mRNA expression. **G.** Schema of folic acid experiment. Serum BUN (**H**), serum creatinine (**I**), and tubular injury scoring after folic acid (24 hours after 250mg/kg IP) (**J**) in iNeph QPRT mice (n=23) and control littermates (n=27). **K.** Representative histology after folic acid iNephQPRT mice and controls. Dotted arrows represent dilated tubules with intratubular casts. Solid arrows represent necrotic cells and denuded tubules. * = p<0.05, ** = p<0.01, *** = p<0.001, **** = p<0.0001.

We next sought to identify transcription factor(s) through which PGC1α co-activates QPRT expression (**Fig 5A**). We first searched the ENCODE Transcription Factor Targets dataset for transcription factors that bind the QPRT promoter based on published ChIP-seq data. ^34^ We next compared that list with known transcription factors that interact with PGC1α in the Human Reference Protein Interactome Mapping Project^35, 36^ and Biogrid^37^ (**Supplemental Table S2**) The intersection identified RXRA and HNF4α as primary candidate transcription factors mediating PGC1α upregulation of *QPRT* gene expression. (**Fig 5B**). Since no RXRA loss of function mutations have been described to cause any renal phenotype in humans based on Clinvar analysis,^38^ and HNF4α loss of function mutations in humans can cause a renal tubulopathy phenotype, we chose to focus on HNF4α.^39^

**Figure 5:**
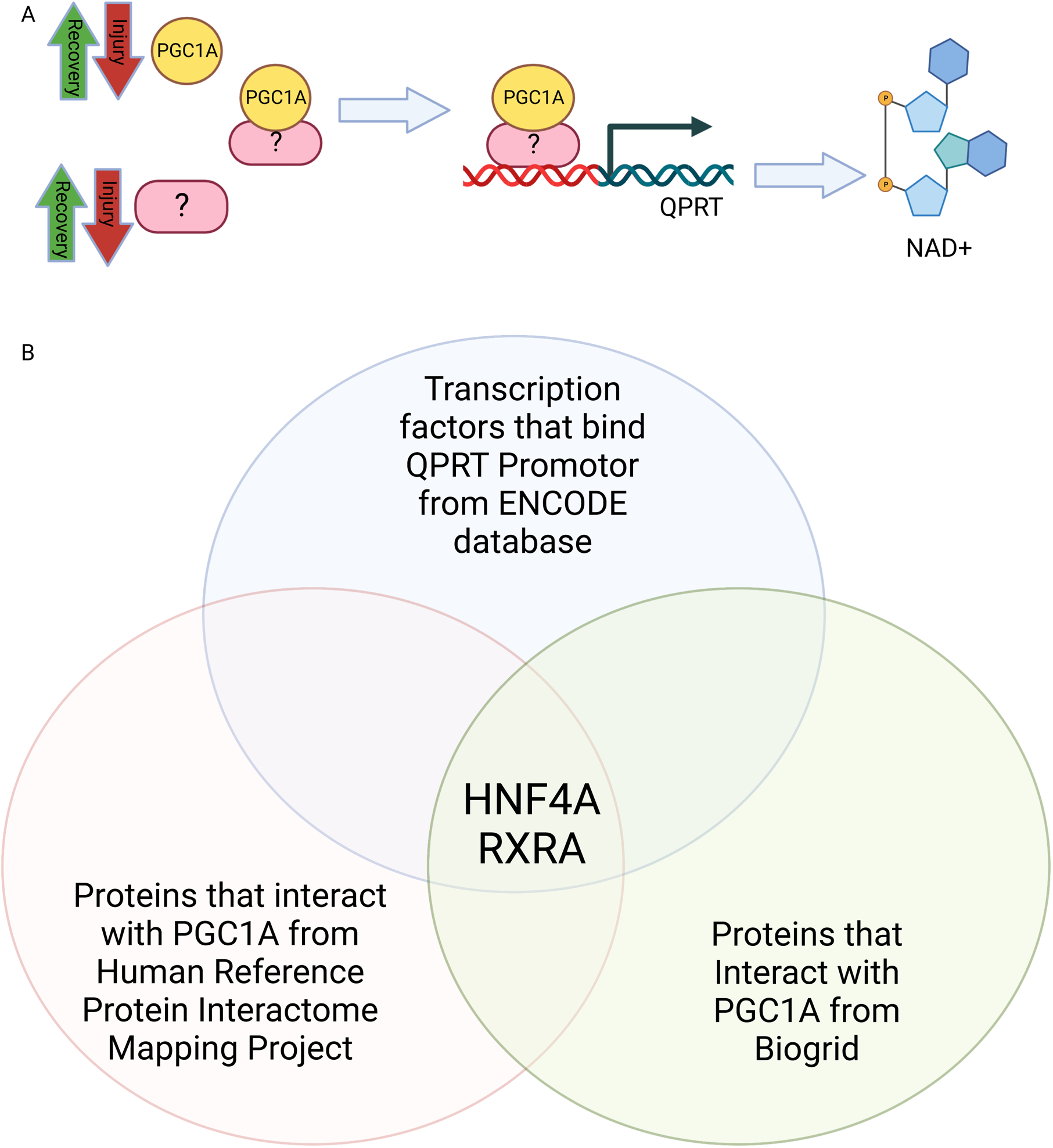
Bioinformatic analysis proposed HNF4α as a transcription factor linking PGC1α and QPRT. **A.** Schematic representing role of unknown transcription factor linking PGC1α and QPRT. **B.** Venn diagram comparing transcription factors known to bind QPRT with transcription factors activated by PGC1α.

Publicly available scRNASeq data of human kidney show that whereas PGC1α is ubiquitously expressed throughout human renal tubule, HNF4α is localized to the proximal tubule (**Fig 6A**).^40^ Likewise, enzymes of de novo NAD+ biosynthesis, including QPRT, localize to the proximal tubule, while enzymes from independent NAD+ biosynthetic pathways do not, consistent with the possibility that HNF4α may be responsible for PGC1α -dependent induction of de novo enzyme gene expression (**Fig 6C**). We therefore sought to determine if expression of HNF4α and QPRT were strongly linked in our models. In kidneys from wild type mice that received cisplatin, HNF4α expression correlated with QPRT expression (**Fig 6C**).

**Figure 6:**
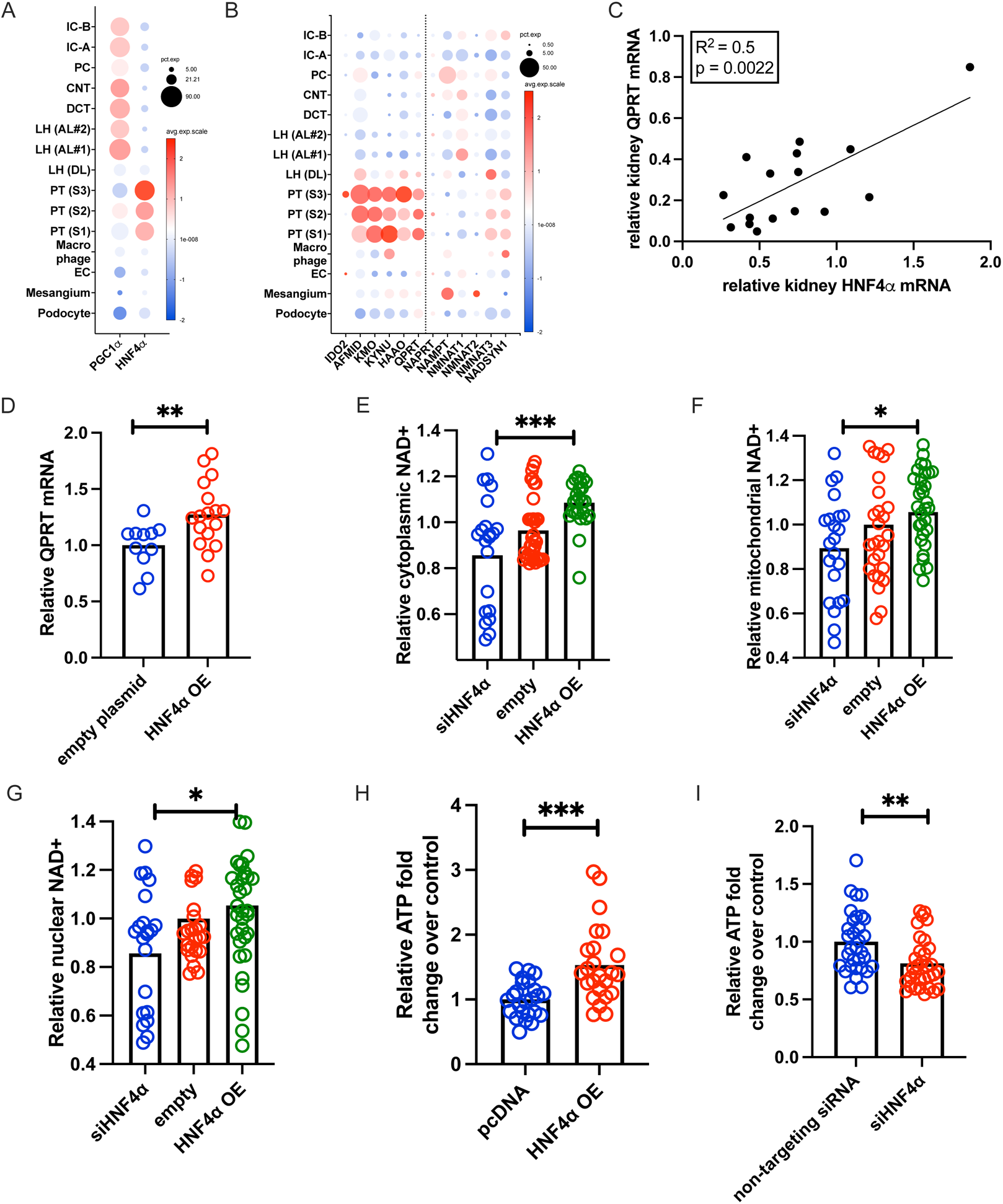
HNF4α activity mirrors QPRT activity. **A.** In human kidney snRNASeq, PGC1α is ubiquitously expressed, but HNF4α localizes to the proximal tubule. **B.** De novo NAD+ biosynthesis and QPRT also localize to the proximal tubule (left of dotted line), while enzymes of other NAD+ biosynthetic pathways are expressed throughout the kidney. **C.** After cisplatin, the degree of QPRT suppression correlates with HNF4α suppression in mouse kidneys. **D.** Overexpressing HNF4α (OE) increases QPRT expression. HNF4α OE increases NAD+, while siHNF4a decreases NAD+ in the cytoplasmic (**E**) nuclear (**F**), and mitochondrial (**G**) compartments. **H.** HNF4α OE increases ATP, and siHNF4α decreases ATP (**I**). * = p<0.05, ** = p<0.01, *** = p<0.001, **** = p<0.0001.

Overexpression of HNF4α in proximal tubule cells increased QPRT expression (**Fig 6D**). To evaluate HNF4α’s effect on cellular NAD+, we co-transfected NAD+ biosensors with a validated HNF4α expression plasmid or HNF4α siRNA (**Supplemental FigS4A-B**). HNF4α overexpression increased NAD+ while HNF4α siRNA decreased NAD+ in each cellular compartment: cytoplasm (**Fig 6E**), mitochondria (**Fig 6F**), and nucleus (**Fig 6G**). Likewise, HNF4α overexpression increased cellular ATP (**Fig 6H**) while siHNF4α decreased cellular ATP (**Fig 6I**). Therefore, the effects of HNF4α and QPRT gene manipulation on NAD+ and ATP were analogous. The results further demonstrate that distinct genetic interventions to manipulate NAD+ result in concomitant changes in cellular ATP, reinforcing the concept that free NAD+ levels are rate-limiting for ATP production in kidney tubular epithelium.

To confirm the PGC1a-HNF4α -QPRT axis in mouse kidney, we performed HNF4α chromatin immunoprecipitation with 2 different antibodies followed by genomic qPCR (ChIP qPCR) for *QPRT* in wild type mouse kidneys and confirmed that HNF4α binds the *QPRT* locus in mouse kidney (**Fig 7A**). We repeated HNF4α ChIP qPCR for QPRT in kidney homogenates from iNephPGC1α mice and littermate controls. These results verified increased enrichment of HNF4α at *QPRT* with PGC1α overexpression (**Fig 7B**) and that the degree of HNF4α pulldown of *QPRT* chromatin correlated with the extent of PGC1α induction (**Fig 7C**). Overexpression of PGC1α (**Supplemental Fig4C)** in human proximal tubule cells increased QPRT expression in cell-autonomous fashion (**Fig 7D**). Moreover, HNF4α siRNA abrogated the PGC1α-dependent increase in QPRT expression (**Fig 7D**).

**Figure 7:**
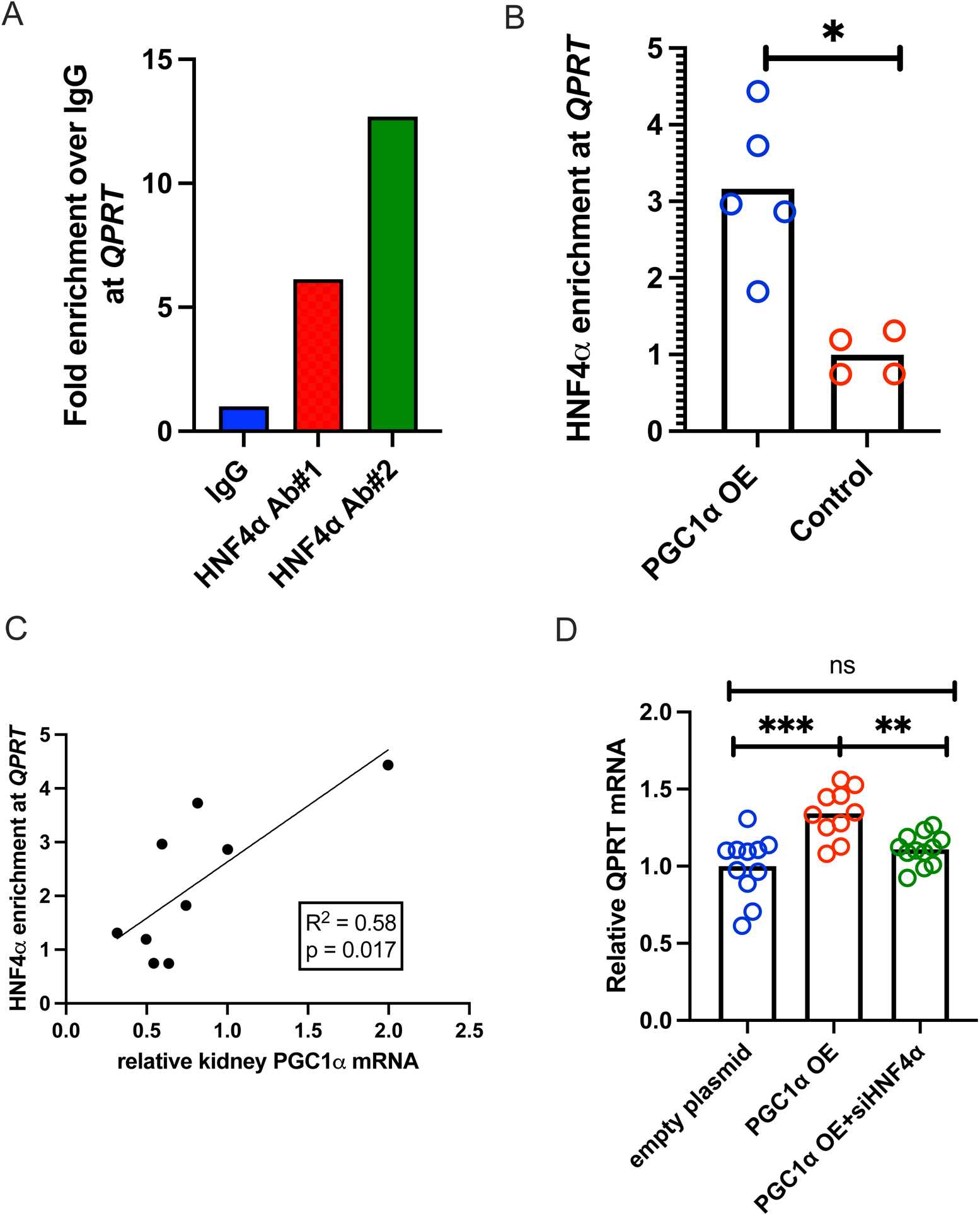
HNF4α modulates renal QPRT, connecting PGC1α to de novo NAD+ biosynthesis. **A.** QPRT qPCR of HNF4α ChIP confirms that HNF4α does enrich at the *QPRT* locus in kidney tissue. **B.** In iNephPGC1α mice, PGC1α overexpression increases HNF4α enrichment at *QPRT*, and (**C**) the level of enrichment correlates with the degree of PGC1α overexpression. **D.** siHNF4α abrogates the PGC1α mediated increase in QPRT expression. * = p<0.05, ** = p<0.01, *** = p<0.001, **** = p<0.0001.

## DISCUSSION

The present studies examined the role of the de novo NAD+ biosynthetic enzyme, QPRT, in experimental acute kidney injury. Previous work linking suppression of this enzyme to AKI was largely limited to post-ischemic models.^7, 8, 10, 11^ The present results advance this concept by evaluating QPRT expression in two distinct models of toxic kidney injury. Cisplatin is an antineoplastic agent that leads to cellular apoptosis via pathologic DNA crosslinking.^41^ Folic acid injures kidney cells via acute tubular obstruction from folate crystals^42^ and subsequent generation of oxidative stress and cellular membrane destruction.^43^ The suppression of QPRT in these non-ischemic settings highlights that de novo NAD+ biosynthesis suppression may be a conserved injury response across different classes of stressors. Therefore, QPRT suppression is likely a feature of the multifactorial AKI that is commonly observed in patients. This has been supported by a study in humans showing elevated urine quinolinic acid to tryptophan ratio in critically ill adults with diverse clinical pathologies who subsequently developed AKI.^7^

Further, the prevention of AKI via kidney tubule specific overexpression of QPRT demonstrates that overcoming the AKI-induced suppression of QPRT is sufficient to restore resilience. This highlights that QPRT enzyme abundance becomes rate-limiting for quinolinic acid flux through the de novo pathway. This is further supported by multiple studies showing that restoration of NAD+ via dietary precursors of alternative NAD+ biosynthetic pathways restores AKI resilience in QPRT haploinsufficent mice^7^ and protects against AKI in wild type animals.^9, 12, 14, 42^

Our cellular results highlight that QPRT enzyme abundance is rate-limiting for both NAD+ and ATP production at baseline. The physiological importance of sustained QPRT levels to combat stress is further evidenced by renoprotection in the inducible gain-of-function mouse. Therefore, two broad possibilities can be considered for a pathogenic role of QPRT suppression following acute stress: (1) decreased flux toward NAD+; and (2) increased accumulation of upstream metabolites in the tryptophan pathway. We have previously shown that AKI susceptibility in the QPRT haploinsufficient state can be “orthogonally” rescued by augmenting NAD+ biosynthesis through an alternate pathway.^7^ However, in support of the latter possibility, quinolinic acid is known to accumulate in CKD and ESKD^11, 45–47^ and is a known neurotoxin.^48^ Even more upstream, buildup of other tryptophan products such as kynurenine may have complex effects on immunity and inflammation.^49, 50^ The long-term implications of metabolite accumulation upstream of QPRT merits further investigation.

While NAD+ is necessary for ATP generation, it has other cellular functions that vary based on cellular compartmentalization. In the nucleus, NAD+ influences gene expression via PARP1 and sirtuins; in the cytoplasm, NAD+ enhances glycolytic flux; and in the mitochondria, NAD+ enhances oxidative respiration.^51^ Using compartmentalized NAD+ biosensors, we found that QPRT modulation affects NAD+ in all compartments. On one hand, this illustrates the cellular significance of QPRT suppression during injury; on the other, it suggests that QPRT suppression affects more cellular functions than oxidative phosphorylation to generate ATP. Previous studies have demonstrated that manipulation of nicotinamide mononucleotide adenyltransferase (NMNAT) isoforms can alter NAD+ concentrations in compartment-specific ways.^51, 52^ Our results provide rationale to explore the role of compartment-specific NAD+ biosynthesis in kidney stress resistance.

Finally, the present results provide novel evidence that HNF4α links PGC1α to the tryptophan pathway of NAD+ biosynthesis. While HNF4α has already been described as a powerful regulator of metabolic pathways in the liver and as a critical component of renal tubular cell differentiation, few studies have examined the metabolic role of HNF4α in the adult kidney. Given the growing evidence of renal metabolism contributing to AKI stress resilience^53^ and even development of chronic kidney disease,^54^ additional studies are urgently needed to understand the breadth of HNF4α mediated gene expression in the healthy and injured kidney. This is especially true as recent multi-omics based investigations have identified recovery of HNF4α expression after injury as one of the most pronounced features of “recovered” tubular cells.^26–28^ These findings also highlight the interesting phenomena of renal tubule cell “de-differentiation” that takes place during injury with loss of expression of genes indicative of differentiated proximal tubule cells, activation of growth factor signaling pathways,^55^ and a reversion from oxidative phosphorylation to glycolysis for energy production.^56, 57^ Loss of HNF4α expression is a hallmark of this phenomenon, and re-expression of HNF4α is one of the most prominent markers of cell recovery.^27^ These findings may implicate de novo NAD+ biosynthesis in cellular differentiation, and additional studies may suggest methods of augmenting NAD+ biosynthesis as a mechanism to modulate cellular differentiation both in injury and in developmental models.

The physiologic activities of HNF4α in the healthy kidney need further study. Our results describe a cell-intrinsic transcriptional axis for regulating tryptophan-dependent NAD+ production. Existing HNF4α fl/fl mice will enable exploration of this axis in animals.^58, 59^ Second, NAD+ metabolism exhibits significant inter-organ crosstalk – for example, the liver and kidney may be the only mammalian organs that synthesize NAD+ from tryptophan and export nicotinamide.^60^ Further evidence of cross-talk comes from rare humans with loss-of-function mutations in this tryptophan pathway manifesting with multi-organ developmental lesions, including prominent renal anomalies.^61^ Therefore, dissecting the organ- and tissue-specific contributions of QPRT will be valuable. For example, studying whether hepatic induction of QPRT can defend against kidney stress is important, particularly since the liver is well-appreciated to influence kidney function. Similarly, conditional deletion experiments are needed.

In summary, the present results demonstrate a conserved and physiologically significant phenomenon of QPRT suppression and NAD+ depletion in response to acute kidney stress, and furthermore, identify a transcriptional mechanism for this response involving HNF4α. These results should advance our current understanding of the relation between NAD+ metabolism and kidney health and disease.

