## Supplemental Figures for "HNF4α mediated QPRT expression in the kidney facilitates resilience against acute kidney injury"

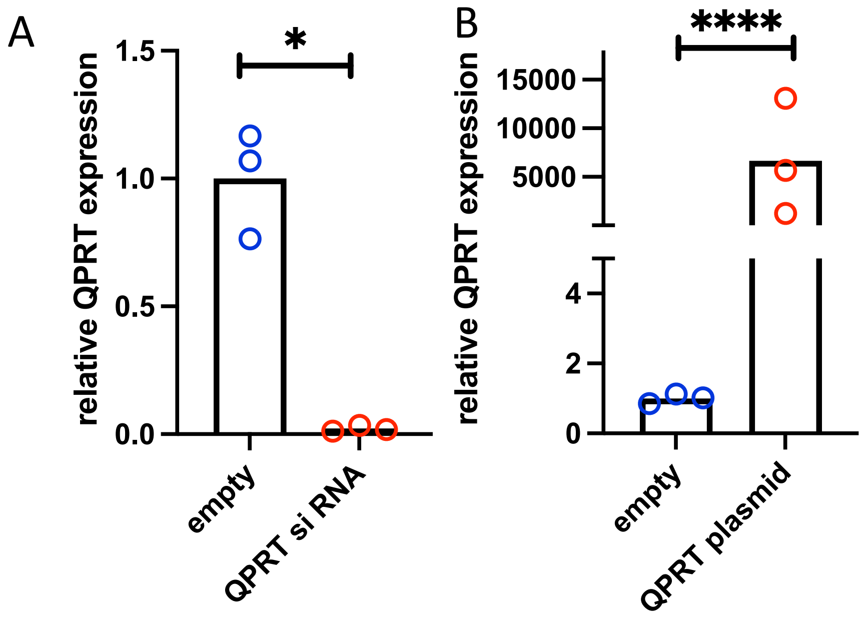


**Figure S1:** Validation of QPRT siRNA (A) and QPRT overexpression plasmid (B) in HK2 cells.

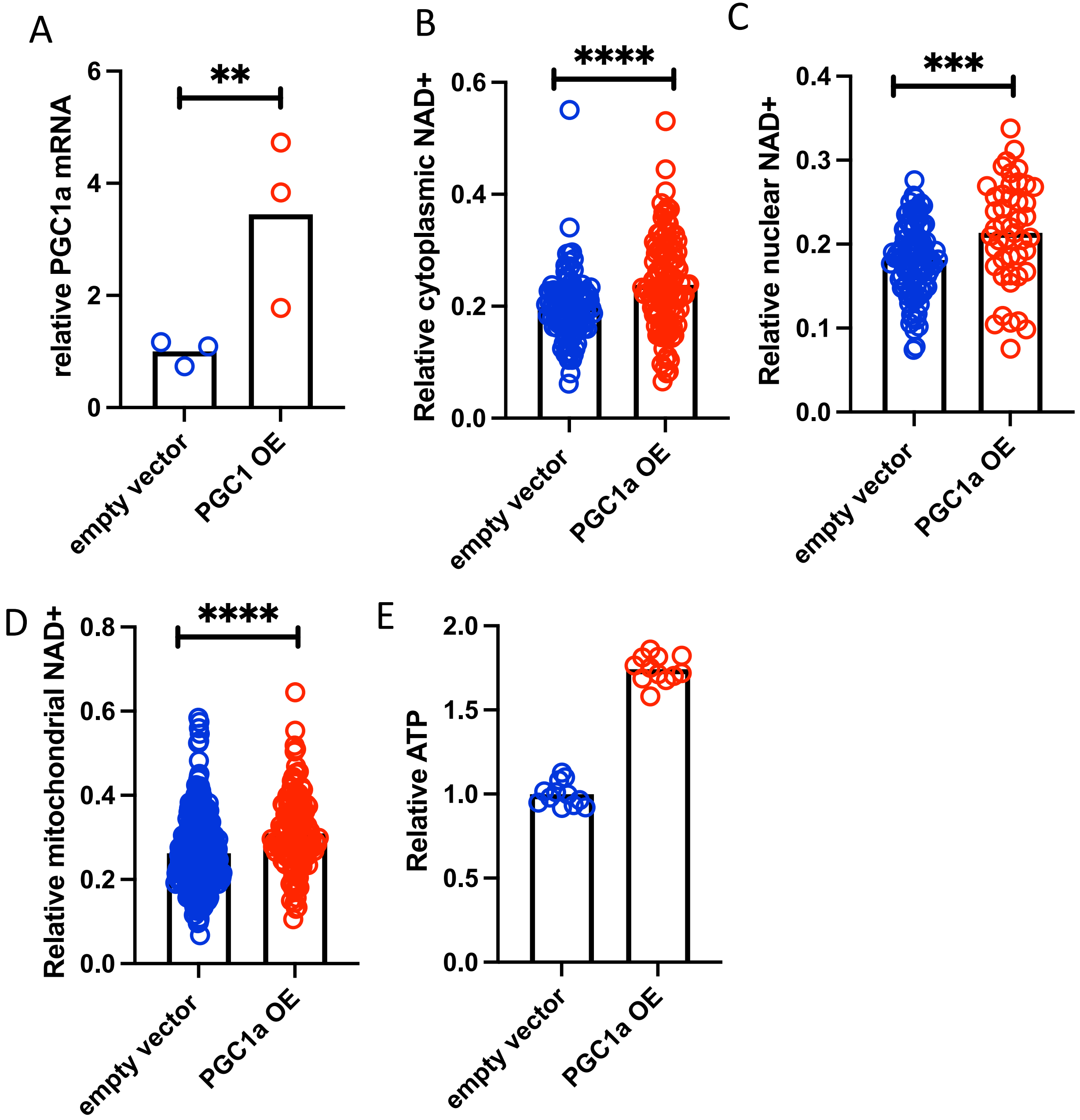


**Figure S2:** A) Validation of PGC1α overexpression in an LLC-PK1 cell line after transduction of Lentivirus carrying PGC1α OE plasmid. PGC1α OE increases NAD+ in the cytoplasm (B), nucleus (C), and mitochondria (D). PGC1α overexpression increases cellular ATP (E).

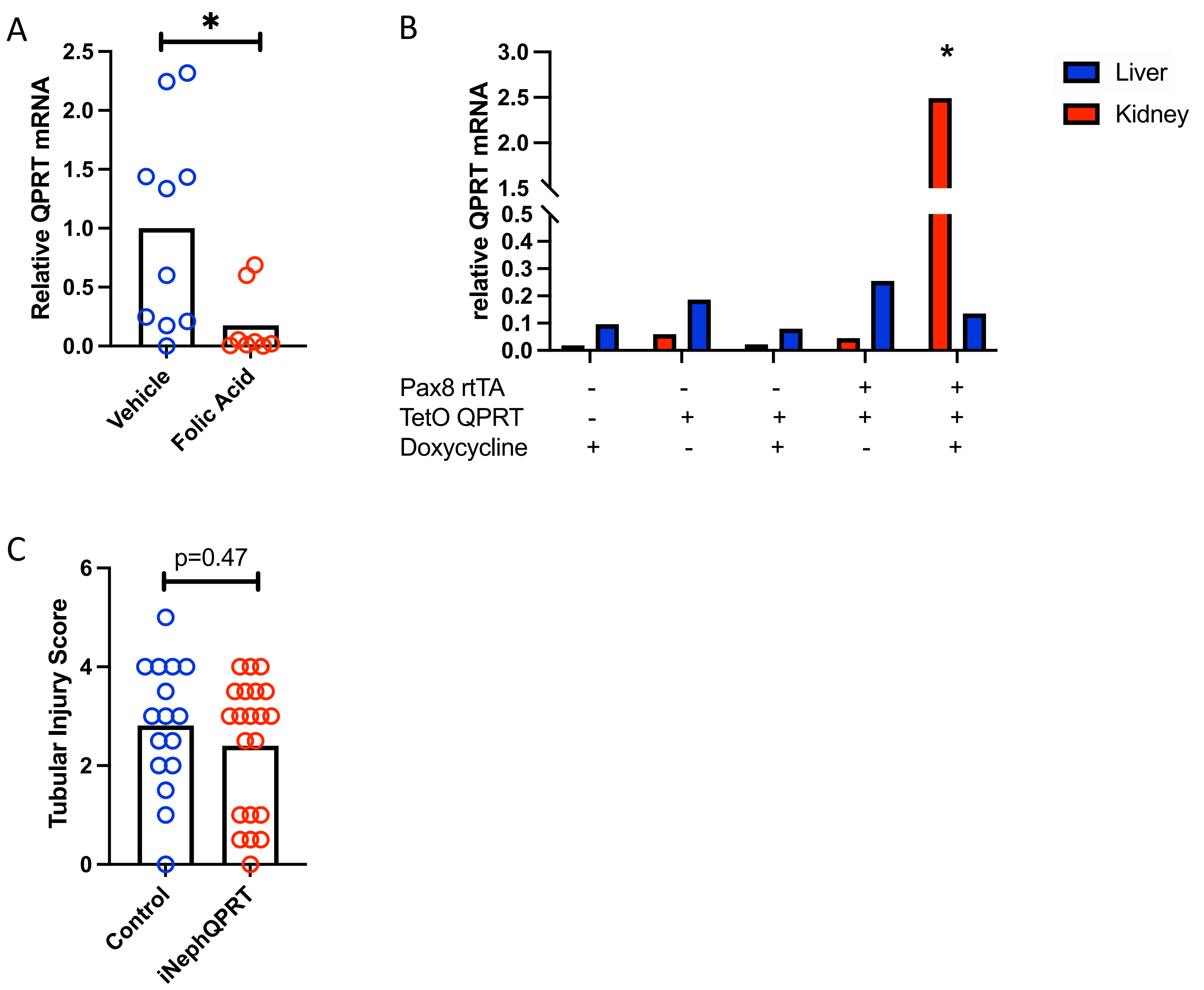


**Figure S3: A.** Folic acid induced AKI leads to QPRT suppression in kidney. **B.** The iNephQPRT mouse exhibits non-leaky overexpression of QPRT in the kidney only when both transgenes are present with the addition of doxycycline. **C.** There is no difference in histological scoring in iNephQPRT mice compared to controls after cisplatin.

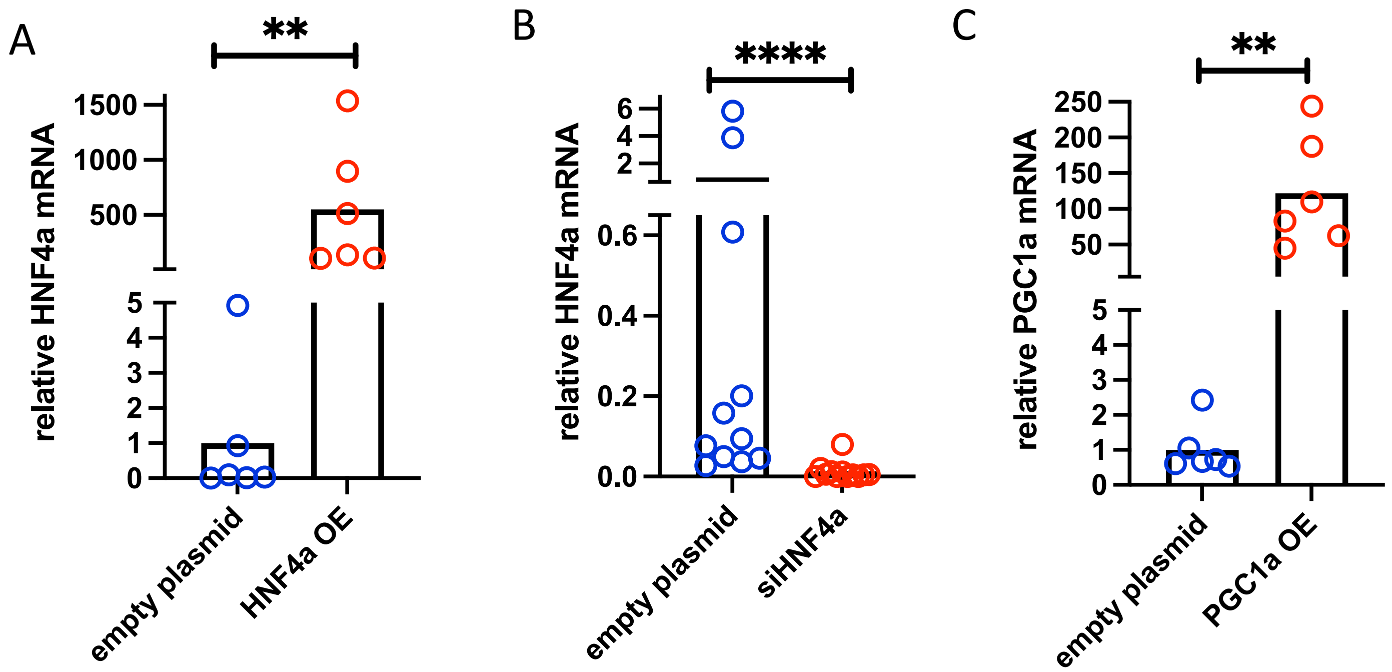


**Figure S4: A-B.** Validation of an HNF4α overexpression plasmid and siHNF4α in HK2 cells. **C.** Validation of PGC1α overexpression plasmid.
